# HoloMoA: Fast holography and deep-learning-based tool for the novelty detection of mechanism of action of antimicrobial candidates

**DOI:** 10.1101/2025.01.17.633645

**Authors:** Zohreh Sedaghat, Benoît Courbon, Héloise Botrel, Hélène Dugua, Pawel Tulinski, Laethitia Alibaud, Lucia Pagani, Derry Mercer, Cyril Guyard, Christophe Védrine, Sophie Dixneuf

**Affiliations:** BIOASTER, 40 avenue Tony Garnier, 69007 Lyon, France; BIOASTER, 28 Rue du Dr Roux, 75015 Paris, France

**Keywords:** Drug discovery, antimicrobial agents, mechanisms of action, holographic microscopy, deep learning, Escherichia coli

## Abstract

We propose an innovative technology to classify the Mechanism of Action (MoA) of antimicrobials and predict their novelty, called HoloMoA. HoloMoA is a rapid, robust, inexpensive, and versatile tool based on the combination of time-lapse Digital Inline Holographic Microscopy (DIHM) and Deep Learning (DL). In combination with proper image reconstruction, DIHM enables a label-free, time-resolved visualization of bacterial cell morphology and phase map (i.e. refractive index × thickness) to reveal phenotypic responses to antimicrobials, while DL techniques are powerful tools to extract discriminative features from image sequences and classify them. We assessed the performance of HoloMoA technology on Escherichia coli (E. coli) ATCC 25922 treated for up to 2 hours with 22 antibiotic molecules representing 5 functional classes (i.e. Cell Wall synthesis inhibitors, Cell Membrane inhibitors, Protein synthesis inhibitors, DNA and RNA synthesis inhibitors). First, using reconstructed phase images as input to a 3D Convolutional Neural Network classifier, we detected the MoA of known antibiotics with 89% accuracy. Secondly, we showed how our CNN models combined with a Siamese neural network architecture can be used for the novelty assessment of the MoA of a candidate antibiotic. The HoloMoA novelty detection tool succeeded in detecting trimethoprim-sulfamethoxazole (Folic Acid synthesis inhibitors) as belonging to a novel functional class (i.e. different from the 5 aforementioned classes). We demonstrated that combining DHIM and DL gives a promising tool for determining the MoA of new antimicrobial candidates provided that a large image database for known antimicrobials is available.

## INTRODUCTION

Antimicrobial resistance (AMR) was responsible for approximately 1.27 million directly attributable deaths in 2019 and was associated with approximately 4.95 million deaths the same year (1). AMR affects countries in all regions, but its effect is exacerbated in low- and middle-income countries (LMIC), generating significant costs. The World Health Organization (WHO) ranked AMR as one of the top global public health threats and pointed out the lack of candidates in the antibiotics pipeline and an access crisis (2, 3).

In the context of the urgent need for the discovery of novel antimicrobials, the WHO updated their Bacterial Priority Pathogens List – first published in 2017 (4) – to guide research and development and public health measures for the prevention, control and treatment of AMR (5). This list ranks 15 families of antibiotic-resistant pathogens into three priority-level groups – critical, high and medium – using several criteria such as mortality, resistance trends, treatability, and antibacterial pipeline. The increasing number of Gram-negative pathogens resistant to last-resort antibiotics on this list underlines the need of novel antimicrobial therapeutic candidates. These new candidates should ideally not induce known cross-resistance, and/or belong to a new chemical class, have new target(s) and have new mechanisms of action (MoA) (3). Based on these criteria, WHO recently reported that, among the 13 new antibiotics that were approved since July 2017, 10 belong to existing classes with known resistance mechanisms. As for the current R&D pipeline, although innovative, the WHO considers it insufficient to combat AMR.

The mechanisms of action underlying the existing antibiotics activities include inhibition of DNA replication, RNA synthesis, protein synthesis, cell wall synthesis, membrane functions, and the production of molecules such as folic acid or ATP (6). Regarding new antimicrobial agents, one of the bottlenecks following phenotypic screenings is identifying the MoA. There is no standard procedure to characterize MoA, so it is highly complex, and fine characterization often involves a combination of several technologies (7), making it expensive and time consuming. A fast, robust, inexpensive and versatile tool, capable of detecting MoA novelty or even the main target of antimicrobial candidates would be very useful to ease the process of antibiotic discovery.

Traditional methods for early-stage MoA classification are labor-intensive and costly. For example, macromolecular synthesis assays (MMS) are an ensemble of 5 assays which involve the incorporation of radiolabeled precursors into macromolecules of the different biosynthesis pathways – peptidoglycan (cell wall), DNA (replication), RNA (transcription), proteins (translation), fatty acids – to identify which pathway(s) is(are) inhibited (8). These assays suffer from low resolution, low accuracy, low throughput and show limitations for the identification of new MoAs. The current need is for an alternative assay that would be cheaper, safer by not requiring the use of radiolabeled precursors, capable of processing not only antibiotic molecules but also complex therapeutic solutions such as drug combinations, and capable of detecting novel MoA.

Recent methods use phenotypic assays that could be applied to library screening campaigns (9), including dynamic metabolome profiling using time-of-flight mass spectrometry (10), transcriptomic (11) and proteomic (12) profiling. Optical methods and in particular imaging methods based on various microscopies are also very attractive, as the accessibility of instrumentation and protocols as well as access to single bacterial cell resolution contribute to a gain in speed and cost (13–17). MoA deduction is based on morphological changes induced by different functional classes of antibiotics on a sensitive strain of a targeted bacterial species (18, 19). Among these methods, bacterial cytological profiling (BCP) is based on fluorescence labeling of DNA and cell wall followed by morphological analysis of each label after a specified period of treatment, either using a confocal microscope (13) or a more common epifluorescence microscope (15). A more recent implementation of label-based fluorescence imaging uses non-cytotoxic labels to exploit the dynamic aspect of morphological changes and increase the resolution and accuracy of classification (16).

Label-free microscopy represents a further step forward in terms of simplification, cost reduction, safety and ecological considerations. No label means that live bacterial cells can be monitored during incubation with the antimicrobial agent, with the guarantee that nothing but the antimicrobial agent will modify the behavior of the cell. Phase contrast microscopy, for example, has been proposed for high-throughput, time-resolved, label-free screening of bacterial morphology to reveal phenotypic responses to antibiotics (17). In the present work, we propose to evaluate an alternative label-free, robust, low-cost and versatile technology for classifying the MoA of an antibacterial and detecting its novelty. The proposed technology, named HoloMoA, combines time-lapse digital inline holographic microscopy (DIHM) and deep learning (DL) analyses. DIHM is the simplest implementation of digital holographic microscopy (20); it enables revealing, in a non-invasive manner, absorption and phase modulations that light undergoes when probing the quasi-transparent bacteria. At 100× magnification, the phase image of a bacterium translates to “thickness × refractive index” heterogeneity within the bacterial cell, without any fixation nor staining. Here, we adapted and evolved an experimental approach, including devices and prototypes, that had been set up during a previous diagnostic DIHM study, to perform antibiotic susceptibility testing of clinical *E. coli* isolates (21–23).

Due to the abundance and high complexity of bacterial time-lapse images, their processing as well as the identification of the antimicrobial MoA must be integrated within an automated pipeline. Increasingly sophisticated statistical methods have been developed to perform such classification. Recent breakthroughs in Machine Learning (ML) and Deep Learning (DL) showed great potential due to their ability to handle large image datasets and extract fine phenotypic information, sometimes not visible to the human eye, to predict a biological outcome of interest. Researchers have proposed a variety of approaches for the automated analysis of high content images to predict the MoA of drugs (24). Basic pipelines rely on extraction of morphological features (such as cell area, mean intensity, roughness…) from the images. This requires prior knowledge from the analyst and possibly the use of image analysis software programs. The features can be combined to perform dimensionality reduction and distance-based clustering to detect MoA groups. Nonejuie *et al*. applied Principal Component Analysis on 14 features derived from ImageJ software followed by Euclidean distance clustering to highlight 5 MoA groups (13). Martin et al. similarly computed 14 morphological features then applied MANOVA and single-linkage clustering to assess the novelty of the MoA of a drug candidate compared to 37 known antibiotics (15). Meanwhile, Ouyang et al. (16) used features extracted from Fiji software (25) and set up a rule-based algorithm to assign antibiotics into 5 MoA classes. A limitation of this approach is the need for prior expertise to compute the features, the presence of uninformative features susceptible to bring noise, and the inability to capture non-linear dependencies among the features. To address the latter aspect, some researchers applied a supervised ML classifier on top of pre-computed features to predict the MoA class of a drug. Zoffman *et al.* trained a Random Forest (RF) on over 100 features to classify 20 antibiotics among 6 MoA classes (14). Likewise, Zahir *et al.* employed Soft Independent Modelling of Class Analogy (SIMCA) to predict 4 morphological groups (17) while Simm et al. (26) computed 842 features with CellProfiler (27) and trained different ML models (Bayesian Matrix Factorization, Random Forest, Deep Neural Network) to predict bioactivity of new compounds.

To overcome the drawback of depending on pre-computed features, end-to-end DL methods have been developed. They have shown great promise due to their ability to extract autonomously a huge number of discriminative features from unstructured data such as images and combine them to perform classification. 2-dimensional Convolutional Neural Networks (CNN2D) have been the most successful architecture to classify images (28). CNN contains convolutional layers which learn automatically and adaptatively spatial features of the data at different scales, in a robust manner. CNN2D and their extensions have shown good performance for MoA prediction based on eukaryotic cell data. For instance, Kensert et al. (29) used deep CNN2D pretrained on large, generic image datasets and fine-tuned them on a dataset of breast cancer cells exposed to 38 chemical compounds corresponding to 12 MoA and achieved high classification performance. Furthermore, Godinez et al. (30) developed a multiscale CNN (M-CNN) that can analyze an image at 7 different resolutions in parallel and achieved cell morphology classification performance superior to existing deep CNN2D models. These methods are efficient to classify known MoAs but are either not designed to detect novel MoAs or deploy only a basic extension of their algorithm to address this question. A common approach is to measure distances in the feature space with respect to known classes. Still, the feature space may lack generalization to unknown data, especially if it is complex and high-dimensional (31). More specific approaches can be used to address novelty detection, such as One Class Support Vector Machines (32), Auto-Encoders (33) or Siamese Neural Networks (34), but to our knowledge, they have not been used for drug screening yet.

In the present work, we propose a fully automated DL pipeline using CNN architecture for MoA classification, taking dynamic holographic data of *E. coli* exposed to bactericidal and bacteriostatic antibiotics as input. Moreover, our pipeline can be adapted for MoA novelty assessment of a drug candidate (i.e. novelty detector). Our approaches introduce two main original aspects compared to state-of-the-art. First, we included a longitudinal component in our DL architecture as we tracked the structural and morphological changes of the bacteria over time. To consider this temporal dimension, we developed two extensions of the standard CNN architecture: 3-dimensional CNN (CNN3D) (23) and Recurrent CNN (RCNN) (35). Second, we used specific DL architecture to provide a robust, quantitative assessment of the novelty of an antibiotic MoA. Specifically, we integrated CNN3D and RCNN into Siamese Neural networks (SNN), resulting in sCNN3D and sCRNN, trained to predict whether two antibiotics share the same MoA.

## RESULTS

### Study design

We used *E. coli* ATCC 25922 which is recommended by the Clinical and Laboratory Standards Institute (CLSI) as a quality control strain for antimicrobial susceptibility tests (36). The holographic image dataset was built with 22 antibiotics split across 11 chemical subclasses and five functional classes constituting 5 “known” MoA: inhibition of DNA replication, inhibition of RNA synthesis, inhibition of Protein synthesis, inhibition of Cell wall biosynthesis, inhibition of Membrane functions. Trimethoprim-sulfamethoxazole is a combination of Folic acid synthesis inhibitors which was used herein to mimic an antibiotic candidate featuring a “novel” MoA and, hence, test the novelty detector, and represents a 6^th^ MoA class. The minimum inhibitory concentrations (MIC) obtained for all these molecules are listed in TABLE S1. We generated the DIHM image dataset (FIG 1–3) before testing the MoA classification tool (FIG 4–5) and novelty detector (FIG 6–7). A single concentration of each compound was used to construct the final image dataset. This concentration was set at 1xMIC, except for kanamycin, tobramycin, tetracycline and lymecyline, which started showing an effect at 2xMIC, and trimethoprim-sulfamethoxazole at 8xMIC. An untreated control was systematically tested for each new bacterial culture. Bacterial cultures and antibiotic exposures took place in Mueller Hinton broth (MHB), with the exception of colistin which required Cation-Adjusted MHB (CAMHB). Each experiment was performed twice, independently. Molecules as well as their corresponding class, tested concentration and medium used are listed in TABLE 1. Each step of the protocol is described in detail in the Materials and Methods section.

**FIG 1.**
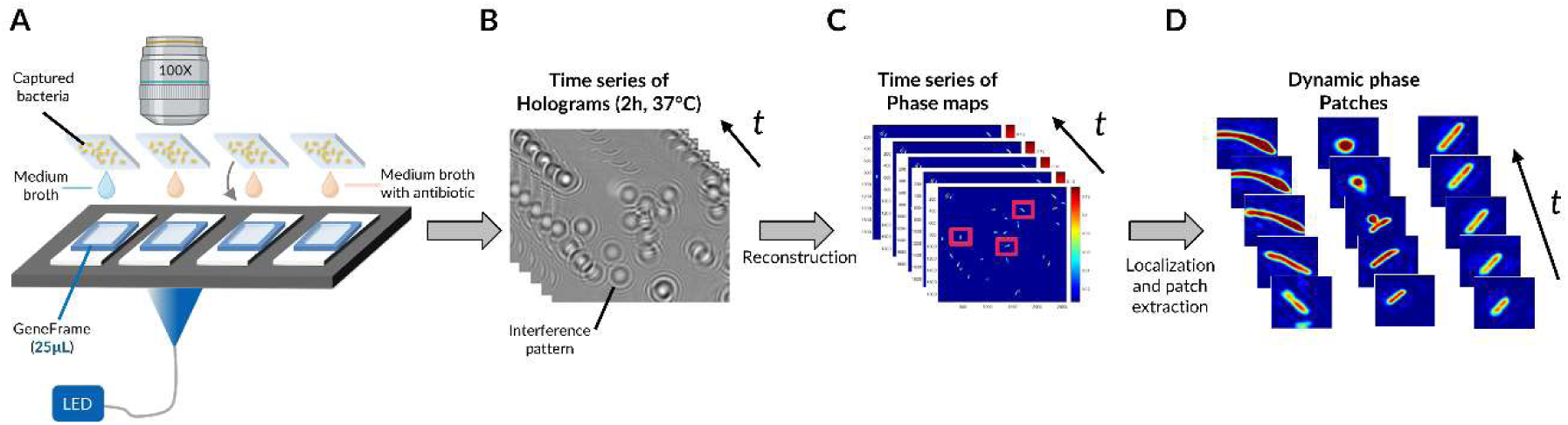
Pipeline for phase image database generation. (**A**) Time-lapse DIHM acquisition for up to 4 conditions (e.g. untreated and treated with 1 molecule at 3 different concentrations). (**B**) Recorded time-lapse holograms for one field of view. (**C**) Reconstructed time-lapse phase maps for the same field of view. (**D**) Dynamic phase map patches resulting from time frame registration and segmentation of bacteria.

**TABLE 1.**
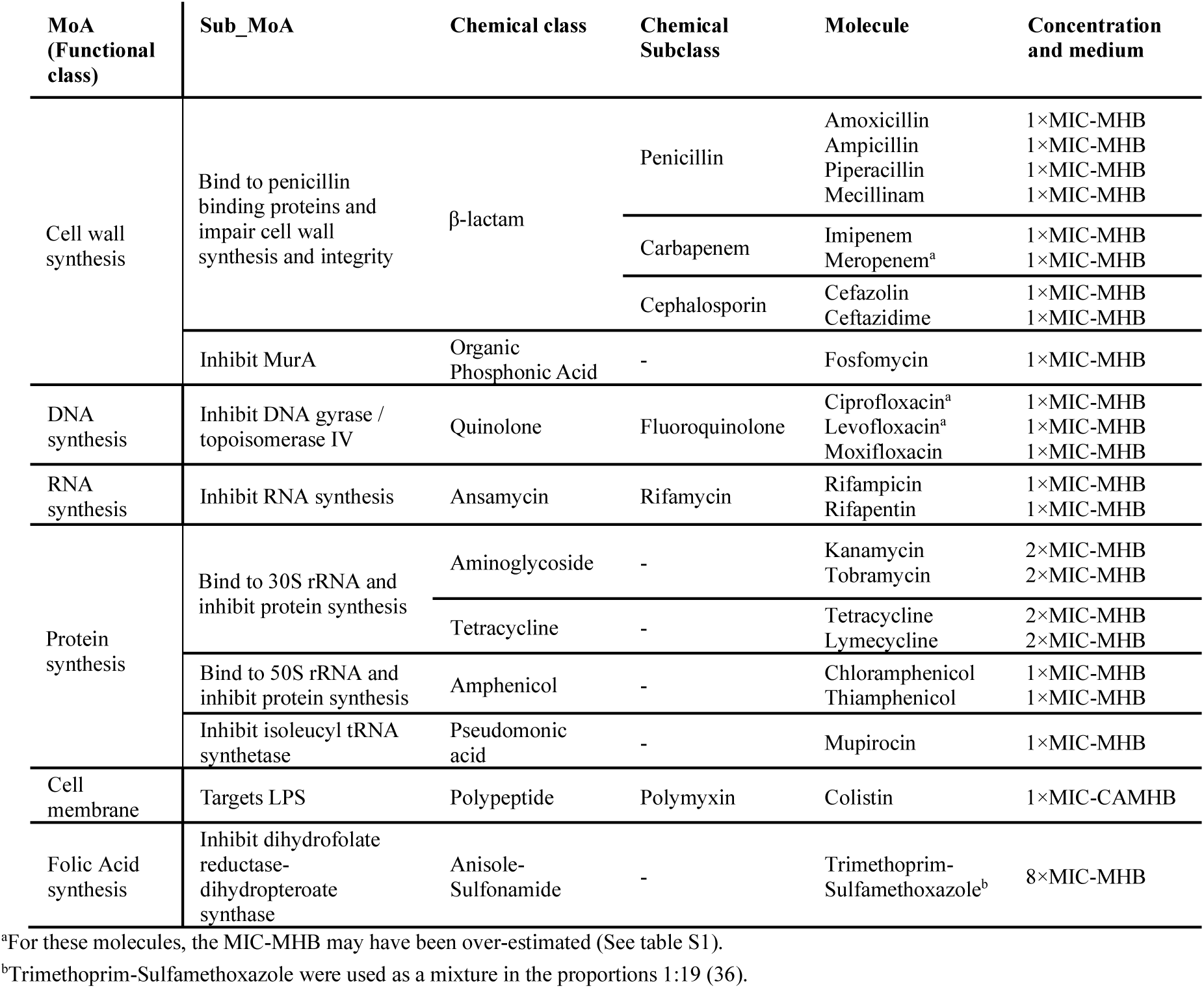
List of antibiotic molecules, corresponding MoA, medium and concentration used for HoloMoA incubations (as a factor of the MIC), number of single-cell dynamic patches.

### Phase image dataset generation

FIG 1 represents the overall analysis pipeline for the phase image database generation. In short, after an in-broth preculture aimed at bringing most bacteria into a metabolically active state, bacterial cells were immobilized via an electrostatic capture and put in contact with antibiotics or not, in a transparent glass chamber suitable for the time-lapse DIHM (FIG 1A and S1). Holograms were acquired every 3 min for 2 h (FIG 1B). Holograms are out-of-focus intensity images featuring interference patterns coding for phase information about the individual diffracting bacteria. Phase images were retrieved following a reconstruction of the acquired holograms, involving an optical-wave propagation algorithm (in the complex plane) based on theory of diffraction of light. Each hologram was reconstructed to obtain in-focus phase images of each field of view and time-point (FIG 1C and S2). Localization of individual bacteria in each field of view allowed us to obtain dynamic patches, each one being a series of temporal images of a single bacterium (FIG 1D).

After having applied this pipeline to the molecules listed in TABLE 1, we obtained a dataset of single bacterial dynamic patches. These patches were preprocessed to remove the ones corresponding to a bacterium fully or partially detached from the coverslip (which could sometimes occur when a bacterium filamented) or to twin image residuals, resulting in unusable images. The preprocessed, final dataset was composed of 1699 single bacteria dynamic patches, each of them being constituted of 40 timeframes of 256×256 pixels vignettes. The dynamic patches split across the 5 “known” MoA classes, the “novel” MoA class, and the “untreated” class (FIG 2). Examples of dynamic patches for each studied molecule are available in the supplementary material (FIG S3). Some of the observed phenotypes over 2 h incubation are illustrated in FIG 3. By eye, the division phenotype of untreated bacteria (FIG 3A) can be distinguished from the absence of division in treated bacteria (FIG 3B-F). Among the treated bacteria, more specific phenotypes emerge depending on the class of drug. Molecules of the Cell Wall class induce strong morphological changes such as elongation, filamentation, sometimes swelling, bulging and/or lysis (i.e. beta-lactams), depending on the specific molecule and drug concentration (FIG S3). These morphological changes correlated often with an increase of the phase (i.e. redder colors on the phase images, FIG 3B). Molecules of the DNA class tended to induce an elongation phenotype and increased phase (FIG 3C). Molecules of RNA, Cell membrane and Protein classes caused similar phenotypes, characterized by a fast freeze of the bacterial morphology; the phase evolution is not easily interpretable by eye for these 3 classes (FIG 3D-F).

**FIG 2.**
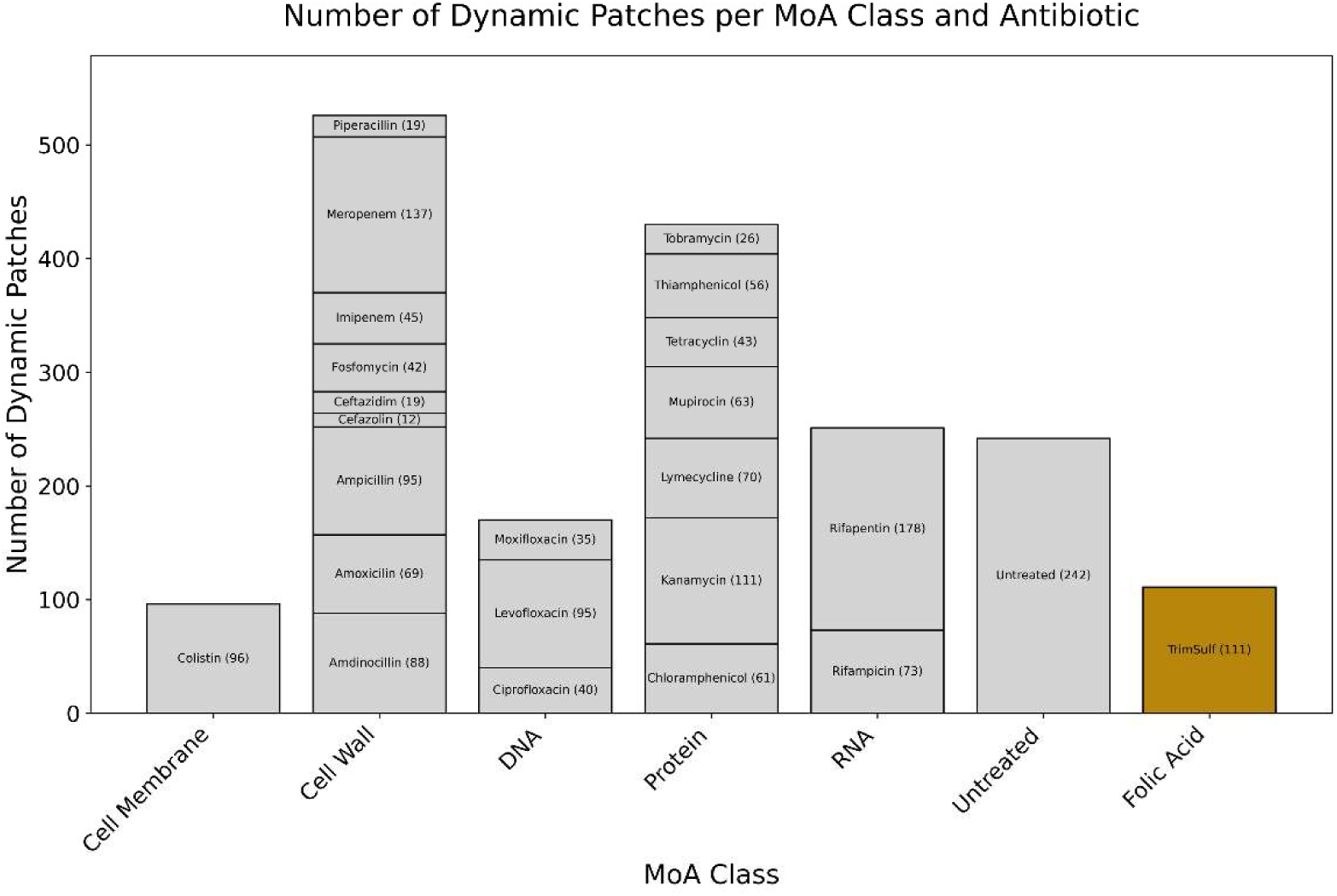
Number of single-bacterium dynamic patches per antibiotic (post preprocessing and cleaning) belonging to the 5 “known” MoA classes of this study (i.e. DNA, RNA, Protein, Cell Wall, Membrane) and to the “novel” MoA class (i.e. Folic Acid), as well as to the “untreated” control class.

**FIG 3.**
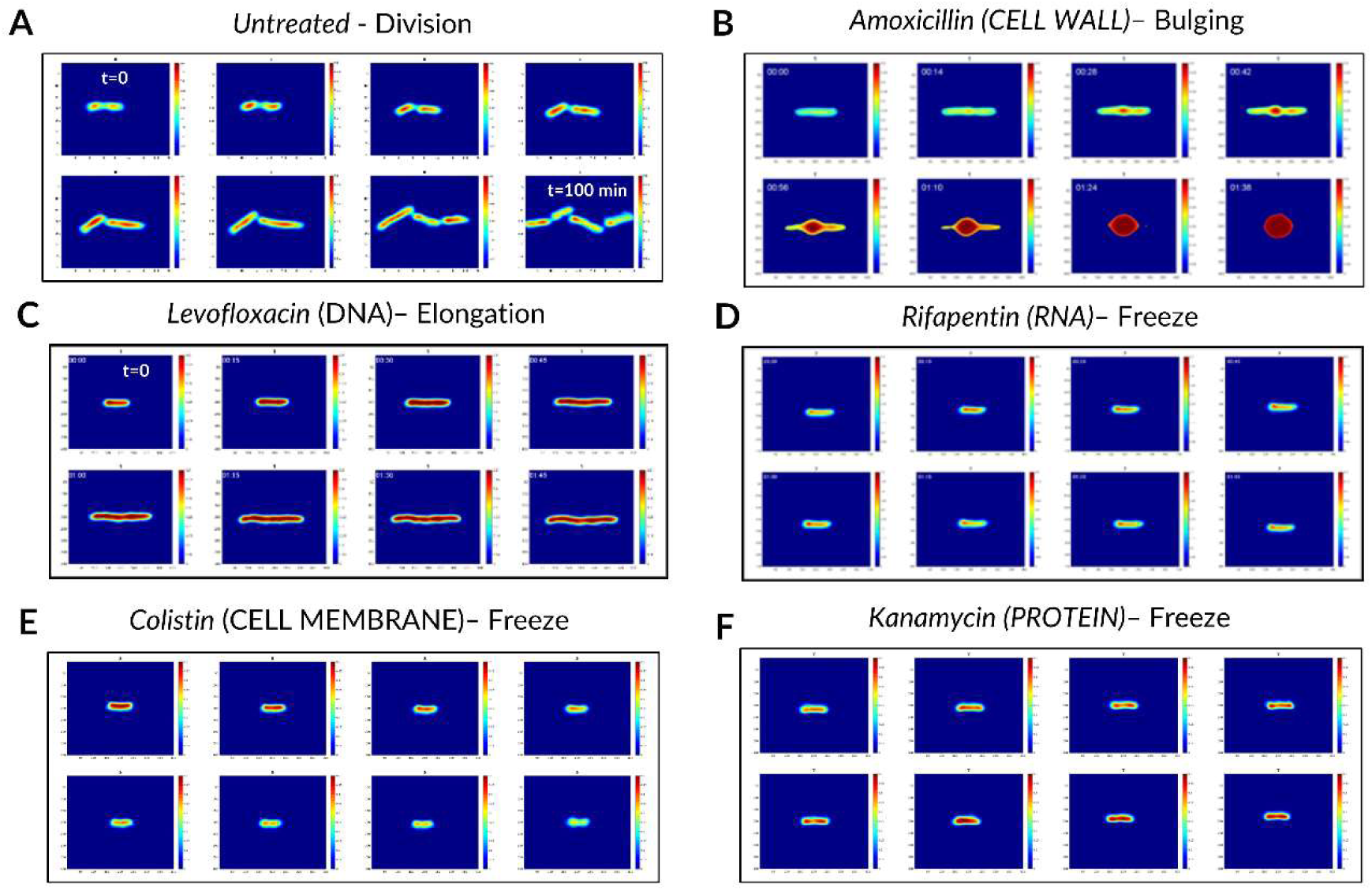
Examples of dynamic phase patches and phenotypes observed over 100 min of incubation without and with antibiotic treatments. (**A**) Division (untreated control). (**B**) Elongation and bulging with strong phase increase. (**C**) Elongation and global phase increase. (**D-E-F**) Frozen morphology and possibly evolving phase change. While images were acquired for 120 min every 3 min, here we show one image every 15 min.

### MoA Classification

We implemented DL classification models to predict the MoA associated with each dynamic patch. We considered 2 main models, described in FIG 4: CNN3D and CRNN. In CNN3D (FIG 4A), the time dimension was analyzed in the same manner as the X and Y space dimensions, using 3D convolutional layers, though with different kernel sizes between time and space. 3D convolutional layers were alternated with pooling layers; classification was achieved through final fully connected dense layers. In CRNN (FIG 4B), the time dimension was treated with a dedicated recurrent layer (Long-Short Term Memory – LSTM, or Gated Recurrent Units - GRU). This layer was incorporated on top of the time-distributed 2D convolutional and pooling layers, followed by the addition of fully connected layers.

**FIG 4.**
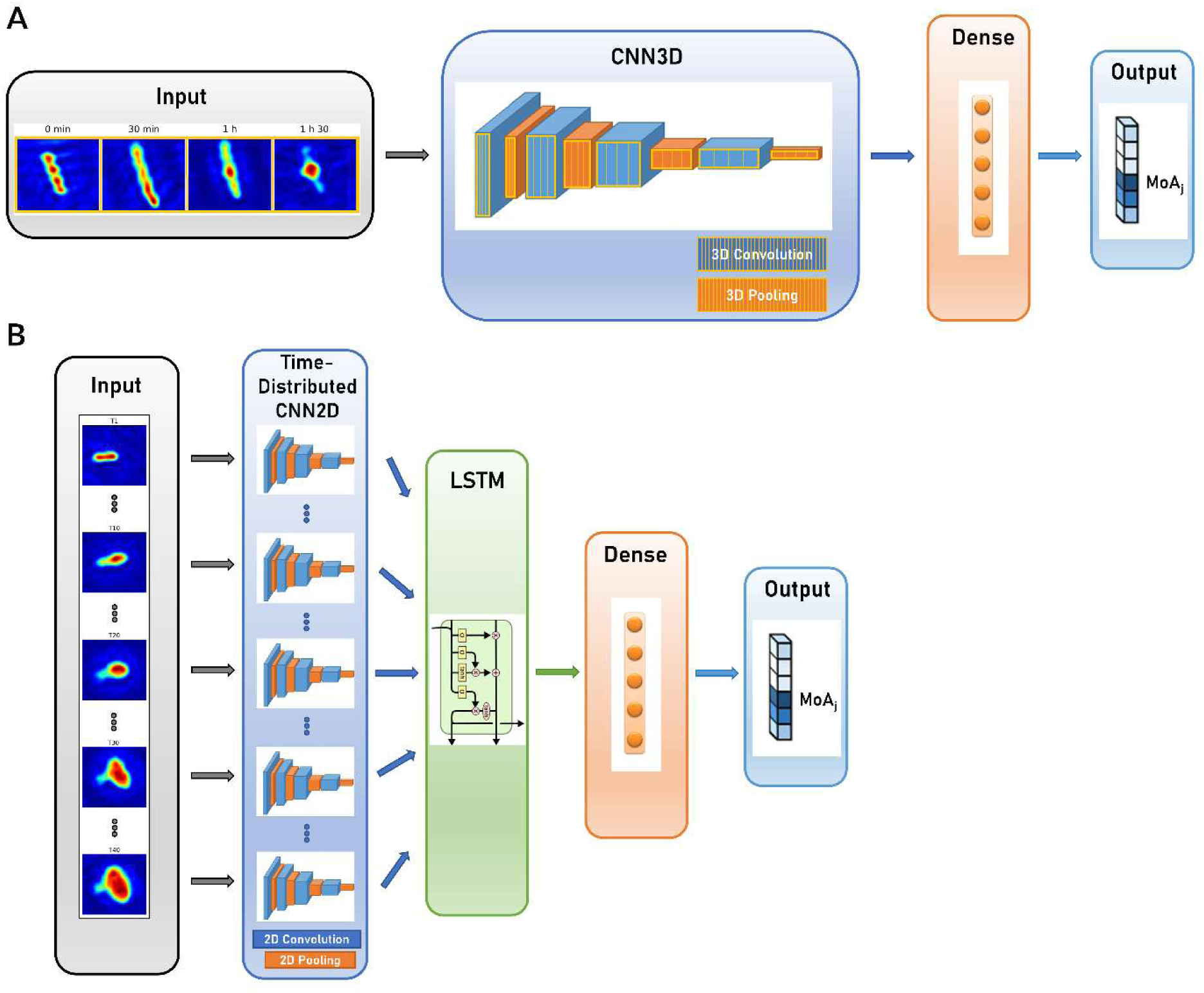
Compared CNN architectures for dynamic patch MoA prediction. (**A**) 3-dimensional Convolutional Neural Network (CNN3D); (**B**) Convolutional Recurrent Neural Network (CRNN).

We used a fast genetic optimization algorithm to optimize the value of hyperparameters such as number of layers, number of units in each layer or kernel sizes (37). We computed the final performance of CNN3D and CRNN models using 10-fold stratified cross-validation. We focused on accuracy (i.e., overall percentage of correct predictions) and F1 score (i.e. the harmonic mean of precision and recall, used to study performance in the case of unbalanced classes). The best model was CNN3D, for which we obtained a dynamic patch classification accuracy of 74% (versus 71% for CRNN) and an F1 score of 73% (versus 68% for CRNN). CNN3D confusion matrix and classification report are shown in FIG 5A and in FIG S4A, respectively. The best performance was achieved for Cell Wall class, with an F1-score of 84%, probably because these molecules induce very specific filamentation or bulging phenotypes (FIG 3A). The main misclassification was obtained between RNA and Protein classes (19% of misclassification rate), for which the size and shape of the bacteria seemed frozen (FIG 3D,F).

**FIG 5.**
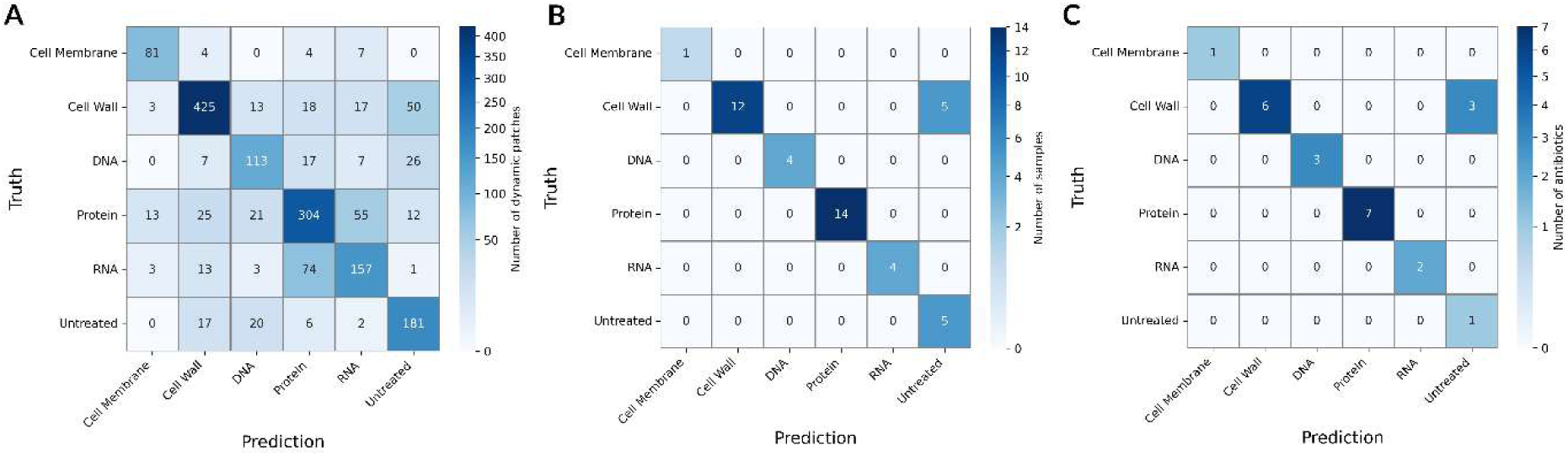
Deep-learning-based (CNN3D) classification results for the untreated class and for 5 known MoA classes: Cell Wall synthesis inhibitors, Cell Membrane function inhibitors, Protein synthesis inhibitors, DNA synthesis inhibitors and RNA synthesis inhibitors. (**A**) Dynamic-patch-level confusion matrix. (**B**) Sample-level confusion matrix. (**C**) Antibiotic-level confusion matrix.

In a second step, we studied the model performance at sample and antibiotic levels by aggregating individual patch predictions. Indeed, in a real-life application, the technology would be applied to assess the MoA from a sample or a collection of samples using the same antibiotic. Hence, to predict the MoA from one sample, we performed majority voting among all its dynamic patch predictions. To predict the MoA of one antibiotic, majority voting was applied to all patch predictions from all replicates associated to the considered antibiotic. This process resulted in a sample-level classification accuracy of 89% and antibiotic-level accuracy of 87% for the CNN3D model. Confusion matrices are shown in FIG 5B and FIG 5C, performance metrics in FIG S4B-C, and classification result per sample and antibiotic in TABLE S2A-B. Model performance was greatly enhanced, which highlights the discriminative power of our pipeline at the slide and sample levels despite inherent experimental and biological variability at single cell level. We observed some confusion between the Cell Wall and untreated classes, due to a subset of samples corresponding to Piperacillin, Ceftazidime and Cefazolin antibiotics. As shown in FIG 2, only a small number of dynamic patches represented these molecules in the dataset, because their important filamentation phenotype resulted in the detachment of many bacteria from the coverslip, which made the hologram reconstruction and segmentation too challenging. This led us to include patches with rather weak filamentation effects, resembling bacteria about to divide in untreated samples. This made it more difficult for the model to learn how to classify them.

### MoA novelty-detection

This work demonstrated the ability of our technology to discriminate known MoA classes using a standard supervised classification scheme. Still, the most needed application for drug discovery is the capacity to predict the novelty of the MoA of an antibiotic candidate compared to a set of known MoA classes. Thus, we proposed a novelty detection framework based on a SNN. The SNN was composed of 2 CNN3D (sCNN3D) or 2 CRNN (sCRNN) with shared weights, trained in parallel and combined with a differencing layer to predict whether 2 dynamic patches correspond to similar (y=0) or different (y=1) MoA classes (FIG 6). During inference phase, dynamic patches from an unknown antibiotic candidate can be compared to the ones from known classes using the trained network. If the candidate exhibits a novel MoA, the network will predict most pairs as different. Our novelty assessment framework is illustrated on FIG S5.

**FIG 6.**
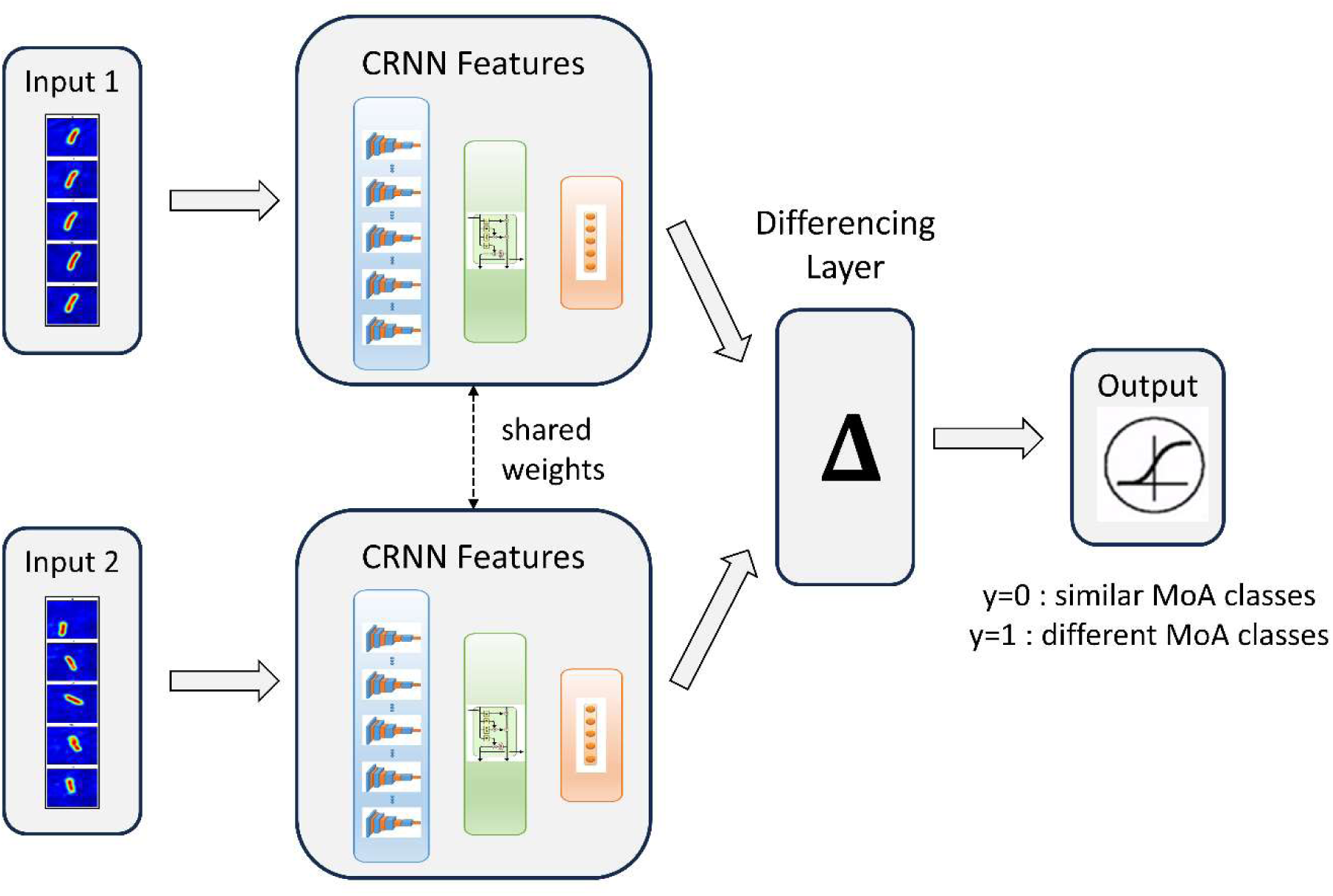
Siamese Convolutional Recurrent Neural Network (sCRNN) architecture for dynamic patches MoA similarity prediction. We call *y* the network prediction.

We designed an experiment to explore the potential of this approach. We defined as known dataset the 5 antibiotic classes and the untreated class, all represented in grey in FIG 2. We considered as an unknown candidate the combination antibiotic trimethoprim-sulfamethoxazole, two molecules belonging to the class of Folic Acid biosynthesis inhibitors. Given that this MoA is different, we expected this candidate to be predicted as novel by our pipeline. We trained our SNN on 70% of the dynamic patches from each known class. We kept 10% of data from these classes in a validation set to compute the intrinsic SNN performance. The remaining 20% were included in a test set, along with data from the unknown candidate that would be used in the inference phase to compute the novelty scores. We precomputed pairs of dynamic patches from the training and validation sets to train and evaluate the SNN. We kept a subset of all possible training pairs to reduce the training time and put more weight on the most important pairs for novelty detection purposes. After subsampling, we obtained 494199 training pairs and 9044 validation pairs.

We trained both sCNN3D and sCRNN models to compare their performances. Our results showed that they were similar (accuracy of 85% and F1 score of 73% for sCRNN versus 84% and 74% for sCNN3D), but sCRNN offered the advantage of reducing training time by half, hence we decided to move forward with this model for the remaining experiment. sCRNN performance metrics and confusion matrix are shown in FIG S6. The model showed good performance at detecting pairs of dynamic patches from different MoAs, as the recall for the class y=0 was 96%. On the other hand, the model frequently misclassified pairs from the same MoA as different, as the recall for the class y=1 was 44%, which is quite understandable given the high biological and experimental variability we observed at the bacteria level.

Then, during the inference phase, we compared all dynamic patches from the test set to the ones in the reference set, composed of the training and validation sets. We aggregated sCRNN predictions per MoA class to compute novelty scores, shown in FIG 7. The novelty scores range between 0 and 1, 0 meaning that the test class exhibits a MoA very similar to the training class, 1 meaning that it is very different. We defined a novelty threshold for each known class (TABLE S3) that corresponds to the novelty score of the class when comparing its training and test data. The novelty threshold is expected to assume a mid-to-low value, depending on the intrinsic variability of the class and the ability of the model to generalize it. If the novelty scores of the unknown class are superior, with a significant confidence margin, to the novelty thresholds of all known classes, the unknown class can be considered as novel. We observed that our experiment validated the proposed framework: trimethoprim-sulfamethoxazole MoA (i.e. Folic Acid) was predicted as novel, since it was assigned novelty scores higher than the corresponding novelty thresholds for all other MoAs (for instance, Protein’s novelty threshold was set at 55%, and Folic Acid scored 68% for this class). As expected, untreated control bacteria samples were confirmed as similar to the reference untreated class.

**FIG 7.**
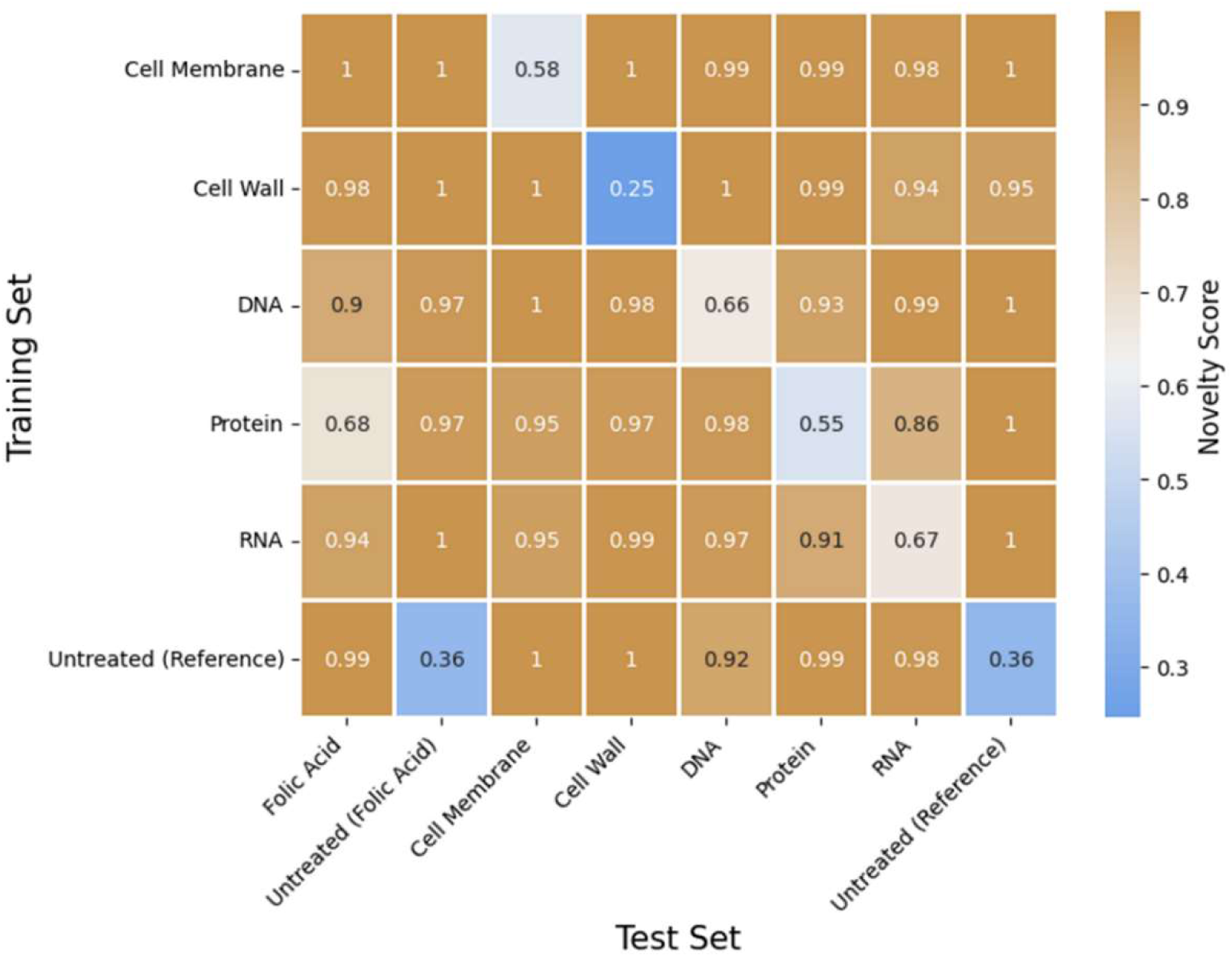
Novelty scores computed for test MoA classes with respect to training (i.e. known) MoA classes, based on pre-trained Deep Learning model (sRCNN).

## DISCUSSION

Herein we have developed a fast, safe and label-free technology to classify antibiotic MoA among 5 functional classes and detect the MoA novelty of an additional antibiotic class using *E. coli* ATCC 25922. This technology, called HoloMoA, combines DIHM and DL. It has proved promising for identifying antibiotics with known MoAs (i.e., whose MoA is in the database) and for detecting the MoA novelty of new antimicrobial candidates. Importantly, HoloMoA takes into account the effect of time on antibiotic action (time-lapse data acquisition). In a context where the determination of a drug MoA remains difficult but important to facilitate preclinical progression, HoloMoA would be more advantageous than the classically used MMS technique.

### Time-lapse DIHM reveals early morphological and phase changes during antibiotic treatment

Time lapse acquisition on single cells allows a focus on rapid antimicrobial effects on the morphology and phase, enabling users to elucidate the antimicrobial MoA in less than 2 h. Changes of bacterial area or mean phase intensity can even be seen as early as 10 min for some molecules (FIG S7-8). Bacterial morphological changes following antimicrobial treatment (19) have already been observed in phase contrast (17), electronic (18) and confocal (13) microscopies. The resolution and contrast obtained with the simple and cost-effective DIHM configuration combined with appropriate hologram reconstruction is close to the more expensive and bulkier confocal microscopy involved for instance in the original BCP technology (13). Both techniques show changes in the shape of the cells. The information retrieved specifically by holographic microscopy is quantitative phase and its 2D heterogeneity within the bacterium area, while confocal images reveal stain-specific parts of the bacterium, typically its cell wall and DNA. We can hypothesize that DL models base their classification on both morphology and phase information (FIG S9). It is interesting to note similarities between phase nodes observed with DIHM and DNA staining observed via fluorescence microscopy, as illustrated in FIG S10. When compared to phase contrast microscopy, DIHM enables the visualization of high phase nods before the actual outer morphology changes, like for instance in the beta-lactam induced bulging phenotype illustrated in FIG S11. Moreover, DIHM does not require precision focusing mechanics as holograms are deliberately acquired out of focus; it is also insensitive to autofluorescence of the sample.

HoloMoA could be coupled to microfluidic approaches for parallelizing many more conditions, automated handling of antibiotic dilutions, delivery to the incubation chambers, and to start imaging from t_0_ of contact of the bacteria with the tested molecules (to avoid the first blind 15 min in the current system). Moreover, the electrostatic capture of the bacteria aiming at their immobilization on the glass substrate to facilitate time-lapse imaging could be replaced by a physical constraint in capillaries on specific microfluidic devices such as a “mother machines” (38), which would prevent bacteria from moving out of the plane when growing or filamenting; this would help increasing the number of usable reconstructed phase images. In addition, DIHM system can be cost-effectively duplicated (See the list of elements in the Materials and Methods section and FIG S1, with motorized stages and the objective as the more costly items) and would be easily adapted to high throughput screening systems, in upright or inverted configurations. Moreover, the required quantity of the tested molecule is minimal (i.e. 25 µL of solution during the treatment phase), which is of importance at the early drug development steps, when available quantities of compounds can be limited.

### DL model performance

Like all AI-based tools for image analysis, the performance of our AI models depends on the size of the dataset, the general rule being “the more the better”. The phenomenon is emphasized when individual data have a high dimensionality, as it is the case for the time-lapse phase image data (i.e. 256 pix × 256 pix × 40 time-points); then datasets will need to include more data examples. The risk of too small a dataset is the lack of generalization. The weak signals are more likely to be ignored due to the lack of examples. It also constrains the models to be carefully tuned to avoid overfitting. In the classification work presented herein, the performance is limited for the Cell Membrane class represented only by colistin, as well as for some molecules of the Cell Wall class, namely Piperacillin, Ceftazidime and Cefazolin, all represented by a rather low number of bacteria (i.e. dynamic patches) compared to the other molecules. For the last 3 molecules, which are beta-lactams, the poor classification performance is worsened by the fact that the patches passing preprocessing criteria are typically the ones showing small filamentation phenotype (by opposition to very long filamentation), therefore resembling the division phenotype of the untreated class. Indeed, filamentous cells detaching even partially from the coverslip over the course of the incubation are eliminated during the automated cleaning of the image dataset. Moreover, the significant cell-to-cell variability cannot be fully avoided, and AI tools may be even more sensitive to it when dealing with time-lapse data. It is likely that the AI models require more examples of variability (i.e. more statistics) when the time dimension is considered (by opposition to endpoint 2D data). Increasing the size and representativity of the image dataset would improve the classification performance.

The highest misclassification rate at single bacterium level was between RNA and Protein synthesis inhibitors (19%) (FIG 5A), both showing almost no morphological changes after treatment (FIG 3 and S3). This may be explained by the fact that protein synthesis, downstream of RNA synthesis, could be blocked as a side effect of RNA synthesis inhibition, which could lead to closely related holographic phenotypes. It is possible the risk of misclassification rate could be reduced by using different antibiotic concentrations or treatment times. Indeed, previous MMS assays testing several antibiotics have shown that these parameters have an impact on the detection of on-target effects, but also of secondary effects or off-target effects (39). Nevertheless, HoloMoA was able to discriminate both these MoAs at the molecule level (FIG 5C and Table S2). An efficient classification at molecule level (FIG 5C and TABLE S2), in addition to the MoA level, suggests that subtle differences between molecules from the same class can be distinguished. This is visible in the Cell wall class where, for instance, imipenem and meropenem (i.e. carbapenems) generated swelling of the bacteria while ceftazidime (i.e. a cephalosporin) induced filamentation (FIG S3), even though both are beta-lactams. This may be explained by the different penicillin binding proteins targeted by each beta-lactam, as reviewed previously (19). These observations show the power of holographic microscopy and suggest that HoloMoA, proposed here as a screening automated method to detect MoA novelty, could also be used to detect even more subtle differences between antimicrobial candidates.

### Focus on the novelty detection feature of HoloMoA

One of the main steps forward of HoloMoA compared to the state-of-the-art lies in the proposal, for the first time to our knowledge, of an automated DL-based pipeline for MoA novelty detection, which could ease the detection of new valuable antimicrobial candidates. Our approach for novelty detection requires relatively high computing resources for siamese neural network training and inference. This is due to the large combinatory arising from the building of pairs to feed the network, associated with the high intrinsic dimensionality of our data. This computing cost prevents us from further optimizing the hyper-parameters of the networks, such as the number of convolutional layers or the subsampling weights. Some solutions can be explored to mitigate this computing cost, such as increasing the subsampling rate or applying a time-space subsampling of dynamic patches themselves. Otherwise, lighter DL architectures could be explored for novelty detection such as the use of convolutional auto-encoders (33), which would reconstruct data from novel antibiotic candidates with a higher loss than data from known antibiotics on which the auto-encoder would have been trained. Moreover, we observe that the MoA classes with lowest statistics (Cell Membrane, RNA, DNA) get the highest novelty thresholds (TABLE S3), due to the lack of generalization of the siamese network. We anticipate that increasing the size and representativity of the image dataset would improve the reliability of our AI tools.

### The use of a wild-type *E. coli* strain

*E. coli* ATCC 25922, which is a CLSI reference strain for antibiotic susceptibility testing, was chosen as the target pathogen for antimicrobial MoA detection and MoA novelty assessment. Its bacillus shape made it a perfect candidate for state-of-the-art microscopy-based examples such as BCP, as it is likely to show more significant morphological changes than a coccoid bacterium like *Staphylococcus aureus* (40–42) or *Neisseria gonorrhoeae*. However, in contrast to most described examples of phenotypic changes caused to bacteria by antibiotics, we preferred using a wild-type strain instead of an organism that has been genetically modified to become “hyperpermeable” to large antibiotic molecules. The hyperpermeable phenotype may alter the morphological changes or even the bacterial response caused by the antibiotics and the relevance of testing Gram positive-only antibiotics against a Gram-negative bacterium may be questionable.

### Summary on improvements and perspective

Among technological improvements foreseen for HoloMoA, we will explore improved immobilization (in the plane) and increased throughput to increase the statistics of image data feeding the AI. We will also test simplification of the image analysis pipeline, using less time-points, full-field images rather than single-bacteria patches, and even go much further and not reconstruct at all the holograms (i.e. use them as is). While bacterial synchronization is not trivial, reinforcing the bacterial preculture step with successive in-broth cultures (43) could optimize the number of metabolically active bacteria when the incubation with the antibiotic starts, and therefore reduce the cell-to-cell heterogeneity. Regarding the biological model, the next step would be to increase the number of molecules in the dataset, up to 60 molecules as in Zampieri et al (10), to reinforce the demonstration by feeding the tool with more phenotype examples, more variability, more off or dual-target cases, etc. The novelty detector should also be tested on real novel antimicrobial candidate molecules. In the longer term, the technology could be implemented for other pathogens including the ESKAPE bacteria but also fungi and parasites. Finally, vibrational spectroscopy such as Raman spectroscopy appears as an appealing label-free modality to use in combination with DIHM. Indeed, Raman micro-spectroscopy would provide the complex chemical fingerprint of the treated bacteria (44), which would complement the morphological and phase information rendered by the DIHM modality. Although access to the time-lapse quantitative phase (via HoloMoA) would already bring more information than standard label-free microscopies such as time-lapse phase-contrast, it is very likely that Raman spectroscopy would bring valuable complementary chemical information to help MoA prediction.

To conclude, thanks to a combination of DIHM and DL methods, HoloMoA demonstrated promising performance for antimicrobial MoA classification and novelty detection, which remain important but challenging requirements during antimicrobial discovery and development. We believe that this tool can support antimicrobial drug developers and, thus, contribute to improving the antimicrobial pipeline and tackle the ongoing AMR pandemic.

## MATERIALS AND METHODS

### Strains, media, and antibiotics

*E. coli* ATCC 25922 was purchased from LGC. Antibiotic powders were dissolved in the appropriate solvent according to manufacturers’ instructions and the resulting stocks were aliquoted and stored at −20°C for a maximum of 6 months. Each aliquot was used only once. MIC were determined following CLSI procedures for broth microdilution antimicrobial susceptibility tests (45). Both MHB (70192, Sigma) and CAMHB (212322, BD) were used: MIC values obtained in CAMHB were compared to the CLSI performance standards (36) to validate the antibiotic stock solutions. Values obtained in MHB were used to design experiments for holographic microscopy analyses, except for colistin which was handled in CAMHB only. The antibiotic MICs are listed in TABLE S1.

### Preparation of the bacteria for holographic microscopy analyses

was done as previously (21). Bacteria from glycerol stocks were thawed and sub-cultured on Columbia agar with 5% sheep blood (COS) (43041, bioMérieux) and incubated overnight at 37°C. The obtained plate was kept at 4°C for a maximum of 3 weeks. Before any further experiment, a single isolated colony was streaked on a new COS plate and incubated overnight at 37°C. The next day, 10 mL MHB or CAMHB were inoculated with isolated colonies diluted to the 0.5 McFarland Standard (McF) (Densimat, bioMérieux) and incubated at 37°C, with shaking at 250 rpm for 2h. Post liquid preculture, 2 × 1 mL were centrifuged for 5 min at 2000 x g at room temperature. After removal of the supernatant, pellets were resuspended in 1 mL of 20 mM phosphate buffer (Phosphate buffer 20 mM; NaCl 50mM; pH 7.2) and centrifuged for 5 min at 2000 x g at room temperature. Pellets were resuspended and pooled in 1 mL phosphate buffer. Turbidity was adjusted to 0.5 McF with phosphate buffer. 120 µL of the suspension were deposited on a 170 µm thick aminosilane-coated glass coverslip (1666121, Schott Nexterion) and allowed to sediment for 15 min at room temperature to allow electrostatic capture of the individual bacterial cells on the coverslip. The coverslip was washed 3 times with the phosphate buffer and once with MHB either with or without antibiotic, while taking care the surface never dried. The coverslip was held with the capture side facing down and placed in a 25 µL incubation chamber, so that the captured bacteria remained in contact with the medium and antibiotic (FIG S1B). The chamber was sealed with 10 × 10 × 0.25 mm^3^ of Gene Frame adhesive (AB0576, ThermoFisher Scientific) fixed on a standard glass microscope slide. The number of captured bacteria over the 10 × 10 mm^2^ surface area was estimated to be ~ 1 x 10^5^ cfu, meaning an inoculum of ~ 10^6^ cfu/mL during incubation with the antibiotics in the 25 µL chamber; this inoculum is comparable to the one used during standard MIC measurement by broth micro-dilution (45). Up to four chambers and incubation conditions were prepared from the same culture and analyzed simultaneously.

### Time-lapse holographic microscopy

was carried out as previously (21), with few modifications. We used a purpose-built prototype of a digital inline holographic microscope (FIG S1A). The light source was a fiber-coupled (M35L01 fiber, Thorlabs), high-power LED emitting in the blue wavelength (FCS-0455-000, Mightex). A 45° protected aluminium turning mirror (CCM1-G01, Thorlabs) enabled direction of the light vertically inside the simplified upright microscope. Light probed the sample held in a 4-slide holder (MLS203P10, Thorlabs); a XY motorized translation stage (NRT100 and MTS25/M-Z8, Thorlabs) enabled horizontal displacement of the samples. A coverslip-corrected 100X objective with a numerical aperture of 0.95 (PLFLN100X, Olympus) collected the transmitted and forward-scattered light; it was mounted on a Z motorized stage to adjust focus (MTS25/M-Z8, Thorlabs). The collected light was filtered by a bandpass filter centered at 450 nm with 10 nm full width at half maximum (FBH450-10, Thorlabs). An achromatic lens with 150 mm focal length (LA1433, Thorlabs) acted as tube lens and focused the collected light on a back-illuminated CMOS camera featuring 3088 × 2076 pixels and 2.4µm pixel size (UI-3884LE-M-GL, IDS). The effective magnification was ~86X. The entire holographic microscope fitted within a standard microbiology incubator set to 37°C. Time-lapse holographic imaging was performed for 2 h in an automated manner via a purpose-made Labview interface, at a rate of one hologram every 3 min and 8 fields of view per sample. For each field of view and time-point (tp), the routine consisted in acquiring one hologram when the sample was defocused by Δz = 20±2 µm (FIG S1C), and one background image when the sample was defocused by Δz = 200µm. Each field of view represented 85 × 57 µm^2^ region of the coverslip and enabled monitoring ~10 captured bacteria. Calibration of the coverslip tilt caused a 15 min delay between the acquisition of the first image of the kinetics and the first contact of the bacteria with the antibiotic. In what follows, t=0 refers to the start of the imaging.

### Hologram reconstruction

The hologram reconstruction algorithm (purpose-built, in Matlab) was mainly performed as before (21). In short, as a preliminary step, each acquired hologram was divided by its corresponding background to normalize and correct for invariant patterns linked to parasitic objects on the light path (i.e. typically dust on optical elements). Second, the reconstruction of the phase shift image (corresponding to the object-plane) from each hologram (corresponding to the image-plane) involved 2 main steps schematized in FIG S2: digital refocalization and twin-image removal. As illustrated in FIG S2A, the propagation algorithm based on Rayleigh Sommerfeld diffraction theory (46) enabled reconstructing a digital z-stack from each hologram; the refocused phase map corresponding to the object-plane was retrieved from the z-stack according to a maximum contrast criterium for almost pure phase objects (i.e. low absorbers) (47). The twin-image artefact is intrinsic to the inline configuration of DIHM and disturbs the background of the reconstructed phase map and its subsequent analysis. We implemented a twin-image removal algorithm inspired by the Gerchberg-Saxton algorithm (48) and its numerous variations, in particular Latychevskaia et al (49). As illustrated in FIG S2B, the algorithm consists in a series of alternating back and forward propagations between the hologram-plane and the object-plane, while imposing constraints on the propagating field in each of the two planes. In the hologram-plane, the squared modulus of the propagated field must be equal to the recorded hologram; in the object-plane, absorption must be positive while the phase shift is forced to 0 when it is lower than a certain adaptive threshold. Iteration after iteration, the reconstructed phase is updated and converges towards its exact value (corresponding to a null threshold).

### Dynamic patches generation

Following the reconstruction of the phase shift maps for each field of view and time-point, individual bacteria were localized via an image binarization to obtain dynamic patches (256 pix × 256 pix × 40 tp) centered on each individual bacterium. Given the strong movements and morphological changes of bacteria (i.e. while dividing or under certain antibiotic treatments), we performed the single-bacteria localization and patch generation on the average image of the 40 time-points. Before entering the machine learning pipeline, a rule-based preprocessing was applied to filter and clean the dynamic patches. The rules were derived from experts’ knowledge and exploratory statistics.

First, dynamic patches failing the quality checks were removed. Three criteria were applied to establish the quality of the images. Phase signal ranged between 0 and 0.5. Dynamic patches starting with pure background images were removed as they were due to segmentation issues. An image was considered as background if it fulfilled one of the following criteria: (i) the maximum phase was below a threshold of *0.3*, (ii) the maximum phase was found in the edge of the image and the mean phase was below *0.02*, where the edge corresponds to a band of *10* pixels width. Dynamic patches with background images at the end of the series were removed. This would correspond to a loss of the bacterium (typically a detachment from the glass coverslip). There was one exception: it could correspond to the lysis of the bacteria, and in that case, we kept the dynamic patch because lysis is an actual signature of the MoA of the antimicrobial. In case of lysis, a small amount of bacteria material remains (“footprint” of the bacteria), therefore we considered as lysis a series of background images at the end of the series with a mean intensity above *0.08.* Dynamic patches with too many empty images were removed. Empty images were due to incorrect holographic reconstruction. These could be corrected with interpolation, as described in the next paragraph. However, we decided to limit the correction applied to the images, to preserve the original biological signal. Then, we removed dynamic patches that fulfilled at least one of these criteria: (i) the patches contained more than *10* empty images in total, (ii) the first image was empty, (iii) more than 2 images at the end of the series were empty.

Second, dynamic patches were cleaned by imputing some empty or background images. (i) Empty images at the end of a series were replaced by a copy of the last non-empty image. (ii) Empty or background images in the middle of the series were replaced by a weighted interpolation of the images before and after them:

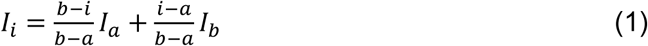

where *I_i_* is the interpolated pixel values of the empty / background image, *I_a_* and *I_b_* the values of the first non-empty / bacteria images before and after it, *i,a,b* the corresponding indices inside the time-series. Interpolation of background images were only performed if only one background image was present between two bacteria images, as it could correspond to a rare, temporary, loss of track of the bacteria.

### Deep-learning-based MoA classification

Two DL-based models were developed for MoA classification. CNN3D (FIG 4A) consisted of a set of convolutional layers associated with maximum pooling layers, a set of fully connected layers combined with dropout layers and finally a classification layer. The convolutional kernels were 3-dimensional and compute convolutions along space and time dimensions. The kernel size was different between space and time dimensions. Convolutional layers activation function was Rectified Linear Unit (“*ReLU*”). Maximum pooling factor was (2,2,2) except after the last convolutional layer where a global maximum pooling operation was performed to combine features and reduce their dimension. Activation functions of fully connected layers were *ReLU*, except for the classification layer which was a fully connected layer with *softmax* activation and a number of units corresponding to the number of MoA classes. CRNN (FIG 4B) consisted of a set of convolutional layers and maximum pooling layers which were time-distributed, a recurrent layer, a set of fully connected layers combined with dropout layers and finally a classification layer. Convolutional kernels were 2-dimensional and squared, activation functions were *ReLU*, maximum pooling factor was (2,2), global maximum pooling was performed after the last convolutional layer. The 2D convolutional networks were time-distributed, meaning that a network was applied to each image in the time-series, but shared the same kernels. Recurrent layers were either Long-Short Term Memory (LSTM) layers or Gated Recurrent Units (GRU) layers. The hyperparameters of the recurrent layers were not optimized compared to the original *tensorflow* implementation. Fully connected, dropout and classifications layers were defined in the same manner as for CNN3D.

Models were trained with Adam optimizer. The model’s loss was categorical cross-entropy. A maximum of 100 epochs was used for training, early stopping was applied with a patience of 15 epochs. Batch size was 8. Final model performance was assessed using a stratified 10-fold cross validation scheme. We optimized the value of various hyperparameters for both models: number of convolutional layers, number of kernels, kernel sizes, number of fully connected layers, number of units per fully connected layers, recurrent layer type, number of units in recurrent layers, dropout rate, learning rate, class weight (meaning whether to re-weight inputs by a factor inversely proportional to the class size). The full list and the ranges that we considered, for each model, are presented in TABLE S4. The last column gives the best value obtained after optimization and was picked for the rest of the workflow. The algorithm we chose for the optimization was the well-known, fast and elitist multi-objective genetic algorithm (37) “Nondominated Sorting Genetic Algorithm II” (NSGA-II) from *Optuna* library (50). Optimization was done using 50 trials and using the model’s loss as metric. Contrary to the final model evaluation, the performance was computed using 4 stratified folds to optimize time. For the same reason, we only used a maximum of 50 epochs with early stopping and a patience of 7 epochs, reducing the batch size to 4.

To compute the classifier prediction at sample and antibiotic levels, we gathered all predictions, corresponding to all dynamic patches, from the 10 validation folds. We aggregated them at the wanted level and we performed a majority vote. We compared the resulting prediction to the true class to compute the model performance metrics at this level. The metric obtained for dynamic patches whose class corresponded to the majority class was called “patch performance” and could be interpreted as a confidence score associated to the sample or antibiotic class prediction.

### Deep-learning-based MoA novelty detection

Novelty detection was based on a siamese neural network (FIG 6) trained on a set of dynamic patches from known MoA classes. The siamese network consisted of two sub-networks sharing the same weights, a differentiation layer and an output layer. The sub-networks were two CNN3D or two CRNN, as described in the previous section with the only difference being that their final classification layer was removed. Their hyperparameters values were the same as the ones obtained after optimization for the classification approach. Each sub-network took as input a dynamic patch. The differentiation layer computed the L1 distance between the outputs of both sub-networks. Finally, the siamese output layer was a fully connected layer of two neurons with *sigmoid* activation, computing a prediction, *y*, which should be equal to 1 if the two input dynamic patches belong to different MoA classes and to 0 if they belong to the same MoA class.

Models were trained with Adam optimizer. The model’s loss was binary cross-entropy. A maximum of 10 epochs was used for training, early stopping was applied with a patience of 3 epochs. The batch size was 4. Model performance was assessed using a stratified validation set containing 10% of input data (135 dynamic patches). The training dataset consisted of 70% of dynamic patches from known classes, i.e. Cell Membrane, Cell Wall, DNA, Protein, RNA and Untreated, corresponding to a total of 1217 dynamic patches. As explained above, the input of the siamese networks was a pair of dynamic patches. To obtain the training pairs set, first we computed all possible pairs, leading to a total of 736291 pairs (578990 corresponding to pairs of different MoA, ie *y=0*, and 157301 corresponding to similar MoA, ie *y=1*) and then we subsampled the dataset so obtained. Subsampling was done for two reasons: to reduce the computing cost, and to allow the model to put more weight on pairs which were the most important to discriminate for the novelty detection task. Specifically, we tried to keep as many pairs as possible from similar MoA, rare MoA classes and MoA classes whose phenotype was difficult to discriminate, as observed during the classification approach. Hence, the subsampling was done according to the following formula:

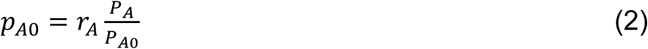

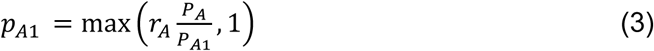

where *p_A0_* is the number of pairs containing dynamic patches from different classes (including class A) after subsampling, *P_A0_* is the number of pairs containing dynamic patches from different classes (including class A) before subsampling, *p_A1_* is the number of pairs containing dynamic patches from the same class A after subsampling, *P_A1_* is the number of pairs containing dynamic patches from the same class A before subsampling, and *R_A_* is a sampling rate defined for each known MoA class, depending on its size and how easy it is easy to discriminate from the others. R_A_ values are listed in TABLE S5.

After subsampling, we obtained 494199 training pairs (336898 corresponding to pairs of different MoAs, i.e. *y=1*, and 157301 corresponding to similar MoA, i.e. *y=0*). We got 9045 validation pairs (7178 corresponding to pairs of different MoA, i.e. *y=1*, and 1867 corresponding to similar MoA, i.e. *y=0*), which were not subsampled. The trained siamese network was then used in inference mode for novelty detection. We computed all possible pairs between dynamic patches from a reference dataset and a test dataset. The reference dataset corresponded to the siamese network training and validation datasets, resulting in 1349 dynamic patches. The test dataset corresponded to the unknown MoA class (Folic Acid inhibitor), the untreated samples associated to this class, and a 20% subset of known classes, resulting in a total of 477 dynamic patches. Therefore, in total we inferred predictions with the siamese network for 643473 pairs. These predictions were aggregated per pair of MoA classes between the reference and the test dataset. For each combination, we computed a novelty score, corresponding to the mean of predictions according to the formula:

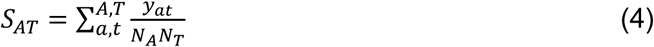

where *y_at_* is the network prediction for a pair of dynamic patches from reference class A and test class A, *N_A_* and *N_T_* are the number of dynamic patches from reference class A and test class *J*.

To assess whether an unknown class corresponded to a novel MoA, the following rule was applied:

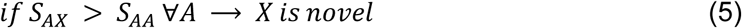

where *X* is the unknown class, *S_AA_* is called the novelty threshold of known class A, and *ε* is a confidence margin.

## SUPPLEMENTARY MATERIAL

Supplemental material. Download HoloMoA_supplementary.pdf

## ACKNOWLEDGMENTS

The authors extend their gratitude to Samuel Bellais, Mélanie Nehlich and Christelle Boisse for their contribution to the standard characterization of some antibiotics, Alexei Novoloaca for helping in a complementary data analysis, Nicolas Sapay for the initial ideas of CNN3D, CRNN and SNN models design, Guillaume Boissy for supporting the Deep Learning acitvity, and Gilles Courtemanche for his encouragement and for initiating efforts around HoloMoA positioning.

